# A Method for Selective ^19^F-Labeling Absent of Probe Sequestration (SLAPS)

**DOI:** 10.1101/2022.06.17.496653

**Authors:** Austin D. Dixon, Jonathan C. Trinidad, Joshua J. Ziarek

## Abstract

Fluorine (^19^F) offers several distinct advantages for biomolecular nuclear magnetic resonance (NMR) spectroscopy such as no background signal, 100% natural abundance, high sensitivity, and a large chemical shift range. Exogenous cysteine-reactive ^19^F-probes have proven especially indispensable for characterizing large, challenging systems that are less amenable to other isotopic labeling strategies such as G protein-coupled receptors (GPCRs). As fluorine linewidths are inherently broad, limiting reactions with offsite cysteines is critical for spectral simplification and accurate deconvolution of component peaks – especially when analyzing systems with intermediate to slow timescale conformational exchange. Here, we uncovered a second source of offsite labeling: non-covalent probe sequestration by detergent micelles. We present a simple four-step protocol for Selective Labeling Absent of Probe Sequestration (SLAPS): physically-disrupt cell membranes in the absence of detergent, incubate membranes with cysteine-reactive ^19^F-probes, remove excess unreacted ^19^F-probe molecules via ultracentrifugation, and finally solubilize in the detergent of choice. SLAPS should be broadly applicable to other lipophilic cysteine-reactive probes and membrane protein classes solubilized in detergent micelles or lipid mimetics.

## Introduction

G protein-coupled receptors (GPCRs) are the largest integral membrane protein class in eukaryotes with over 800 unique members that regulate numerous biological processes including mood, body temperature, taste, and sight, amongst others.^1,2^ They share a conserved architecture of seven transmembrane (TM) alpha-helices that bundle together to form an extracellular orthosteric binding pocket and an intracellular cytosolic cleft.^3^ Ligand binding at the orthosteric pocket induces a conformational change at the intracellular cleft to enable G protein association, guanine nucleotide exchange, and ultimately the intracellular signaling cascade. Termination of GPCR signaling is mediated through ternary complex formation with Arrestin, which activates clathrin-mediated endocytosis for receptor recycling/degradation.^2^ Due to their broad physiological importance and numerous etiological roles, GPCRs are the targets for more than 30% of all therapeutic drugs on the market.^4^ A more nuanced mechanistic understanding of the GPCR activation landscape could dramatically expand their therapeutic value.^5^

Spectroscopic techniques such as fluorescence,^6^ infrared (IR),^7^ electron paramagnetic resonance (EPR),^8^ and nuclear magnetic resonance (NMR)^9^ have revealed many lowly-populated, high energy conformational states that remain invisible to X-ray crystallography and cryo-EM. In particular, the ability of NMR to access motional regimes covering more than 15 orders of magnitude (ps-s) makes it especially attractive for this task, although the challenges associated with uniform incorporation of NMR-active isotopes has somewhat limited its application.^10^ Exogenous cysteine-reactive fluorine (^19^F) probes have proven an effective alternative to uniform labeling^11–16^ owing to their high gyromagnetic ratio (i.e. sensitivity), 100% natural abundance, large chemical shift range, and absence of background signals in biomolecular samples.^17^ Yet, fluorine’s intrinsically-broad linewidths quickly lead to overlapping signals that require deconvolution, and generally prohibits the simultaneous labeling of multiple sites. Offsite ^19^F-probe incorporation is an additional source of signal overlap that is specifically problematic when the target protein contains critical cysteine residues that cannot be mutated.^9,18,19^

In our previous work labeling the neurotensin receptor 1 (NTS1) Class A GPCR with cysteine-reactive ^19^F-probes,^16^ we uncovered a second source of offsite labeling: non-covalent sequestration by detergent micelles. Conventional labeling methods solubilize the receptor in detergent micelles without the prior removal of excess ^19^F-probe molecules. Our liquid-chromatography mass spectrometry (LC-MS) and NMR spectra of a cysteine-less NTS1 construct demonstrate that unreacted ^19^F-probe molecules are sequestered into proteomicelles. Subsequent detergent wash steps or detergent exchange is incapable of complete excess ^19^F-probe removal. We present a simple four-step protocol for Selective Labeling Absent of Probe Sequestration (SLAPS): physically-disrupt cell membranes in the absence of detergent, incubate membranes with cysteine-reactive ^19^F-probes, remove excess unreacted ^19^F-probe molecules via ultracentrifugation, and finally solubilize in detergent of choice.

## Results and Discussion

Several generations of thiol-reactive trifluoromethyl probes have been developed to study GPCR dynamics.^20–22^ 2-bromo-N-(4-(trifluoromethyl)phenyl)acetamide (^19^F-BTFMA) remains one of the most popular probes due to its ability to form a nonreducible thioether bond, along with high chemical shift sensitivity owing to aromatic ring polarizability [Fig. 1(A)].^22^ The majority of ^19^F-GPCR studies conjugate probe to the intracellular tips of TM5,^23^ TM6,^24^ or TM7,^25^ which have proven invaluable for mapping the receptor activation landscape due to their large architectural changes.^26^ In many cases, this requires the introduction of a non-native cysteine residue at the position of interest and the simultaneous mutagenesis of all endogenous solvent-exposed cysteine residues that would lead to offsite labeling. Nonetheless, researchers have noted the presence of offsite ^19^F-labeling in final protein samples.^9,18,19^ These are commonly attributed to the numerous reduced cysteine resides in the transmembrane region, although, few have been verified [Fig. 1(B)].

**Figure 1.**
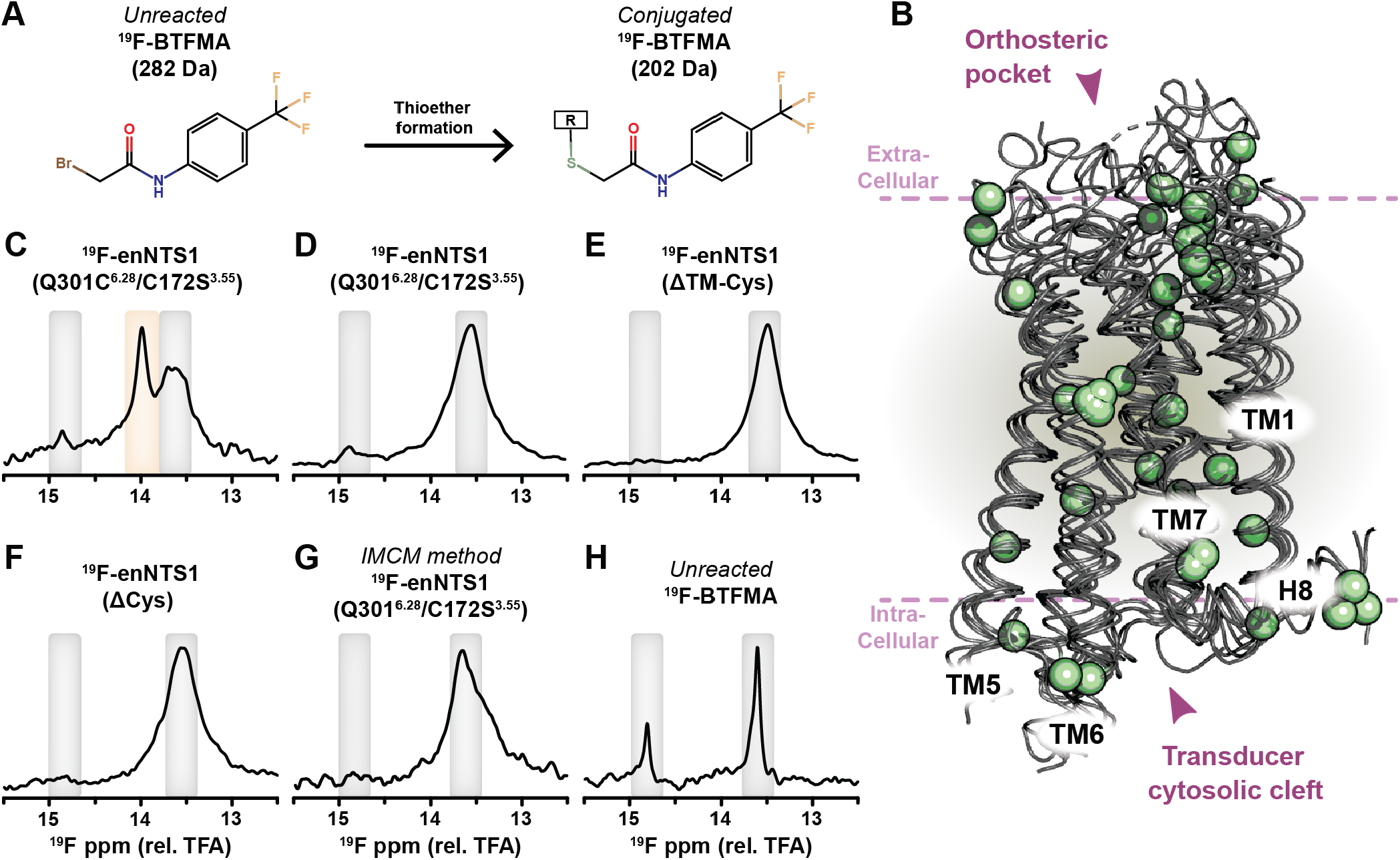
Labeling detergent-solubilized enNTS1 results in offsite ^19^F-BTFMA probe incorporation. (A) Unreacted ^19^F-BTFMA (*left*) conjugates to cysteine residues via thioether bond formation (*right*), adding +202 Da to receptor molecular weight. (B) Overlay of β_1_-adrenergic (PDB 4BVN), β_2_-adrenergic (PDB 2RH1), Adenosine A_2A_ (PDB 4EIY), Rhodopsin (PDB 1U19), and Neurotensin receptor 1 (PDB 4BWB) atomic models illustrates the numerous cysteine residues (green spheres) located throughout the extracellular, transmembrane, and intracellular regions.^33–37^ NMR spectra of (C) ^19^F-enNTS1(Q301C^6.28^/C172S^3.55^), (D) ^19^F-enNTS1(Q301^6.28^/C172S^3.55^), (E) ^19^F-enNTS1(ΔTM-Cys), (F) ^19^F-enNTS1(ΔCys), (G) ^19^F-enNTS1(Q301^6.28^/C172S^3.55^) prepared by IMCM, and (H) unreacted ^19^F-BTFMA solubilized in detergent micelles. ^19^F-chemical shifts are relative to trifluoroacetic acid (TFA). Final sample buffer conditions for all NMR spectra: 20 mM HEPES, 50 mM NaCl, 50 µM TFA, and 0.01% (w/v) LMNG at pH 7.5. All NMR samples were supplemented with 10% (v/v) D_2_O.

Spectroscopic studies require receptors to be isolated from native lipid membranes into detergent micelles for purification and, frequently, analysis. The ^19^F-probes are typically incorporated following detergent solubilization of native lipid membranes, but prior to purification. We applied this strategy to label a thermostabilized Neurotensin receptor 1 variant (enNTS1).^27^ We introduced an exogenous cysteine on TM6 (Q301C^6.28^, Ballesteros-Weinstein nomenclature^28^) and substituted the only solvent-exposed cysteine (C172S^3.55^) to reduce offsite labeling, referring to the final construct as enNTS1(Q301C^6.28^/C172S^3.55^). Briefly, *Escherichia coli* cell pellets containing enNTS1(Q301C^6.28^/C172S^3.55^) were resuspended in aqueous buffer, sonicated, solubilized with 1% (w/v) n-decyl-β-D-maltopyranoside (DM) detergent, and incubated for 1 h with ^19^F-BTFMA. The sample was immobilized on metal-affinity resin for exchange to 2,2-didecylpropane-1,3-bis-β-D-maltopyranoside (LMNG) detergent micelles, then purified by cation exchange and gel filtration. The ^19^F-enNTS1(Q301C^6.28^/C172S^3.55^) 1D ^19^F NMR spectrum contained three resonances at 13.6, 14.0, and 14.8 ppm [Fig. 1(C)]. Many previous ^19^F-GPCR studies reveal that TM6 exchanges between multiple conformations on the ms-s timescale, which would similarly produce a spectrum containing multiple peaks, even when ^19^F-labeled at a single position (i.e. no offsite labeling).^9,13,16^ As a negative control, we engineered ^19^F-enNTS1(Q301^6.28^/C172S^3.55^) with residue 301 reverted to glutamine and repeated the experiment. The spectrum contained two resonances that were present in the ^19^F-enNTS1(Q301C^6.28^/C172S^3.55^) spectrum indicative of offsite labeling [Fig. 1(D)].

We generated two additional cysteine-depleted enNTS1 constructs, enNTS1(ΔTM-Cys) and enNTS1(ΔCys) to identify which cysteine residue was being labeled [Fig. 2(A)]; both constructs included C172S^3.55^ and reverted residue 301^6.28^ to glutamine. enNTS1(Q301^6.28^/C152S^3.35^/C172S^3.55^/C320S^6.47^) eliminates the reduced cysteine residues from the transmembrane region (C152S^3.35^/C320S^6.47^) while enNTS1(ΔCys) is entirely devoid of cysteines (C142S^3.25^/C152S^3.35^/C225S^ECL2^/C320S^6.47^). Both constructs were again ^19^F-labeled following DM detergent solubilization and purified as above. Surprisingly, ^19^F-enNTS1(ΔTM-Cys) and ^19^F-enNTS1(ΔCys) spectra both contained a strong resonance at 13.6 ppm and a weaker one at 14.8 ppm as observed in the other ^19^F-enNTS1 spectra [Fig. 1(C-F)]. Wüthrich and colleagues recently showed that detergent-solubilized receptors were highly reactive and proposed the In-Membrane Chemical Modification (IMCM) approach to reduce offsite labeling.^19^ IMCM exploits the membrane’s natural protection of transmembrane cysteine residues by conjugating the probe following physical disruption of the lipid bilayer *but prior to* detergent solubilization; after probe incubation the receptor is solubilized in detergent and purified. We applied IMCM to our ^19^F-enNTS1(Q301^6.28^/C172S^3.55^) negative control by incubating sonicated membranes with ^19^F-BTFMA for 1 h prior to solubilization in DM micelles. However, we still observed a strong ^19^F-resonance [Fig. 1(G)].

**Figure 2.**
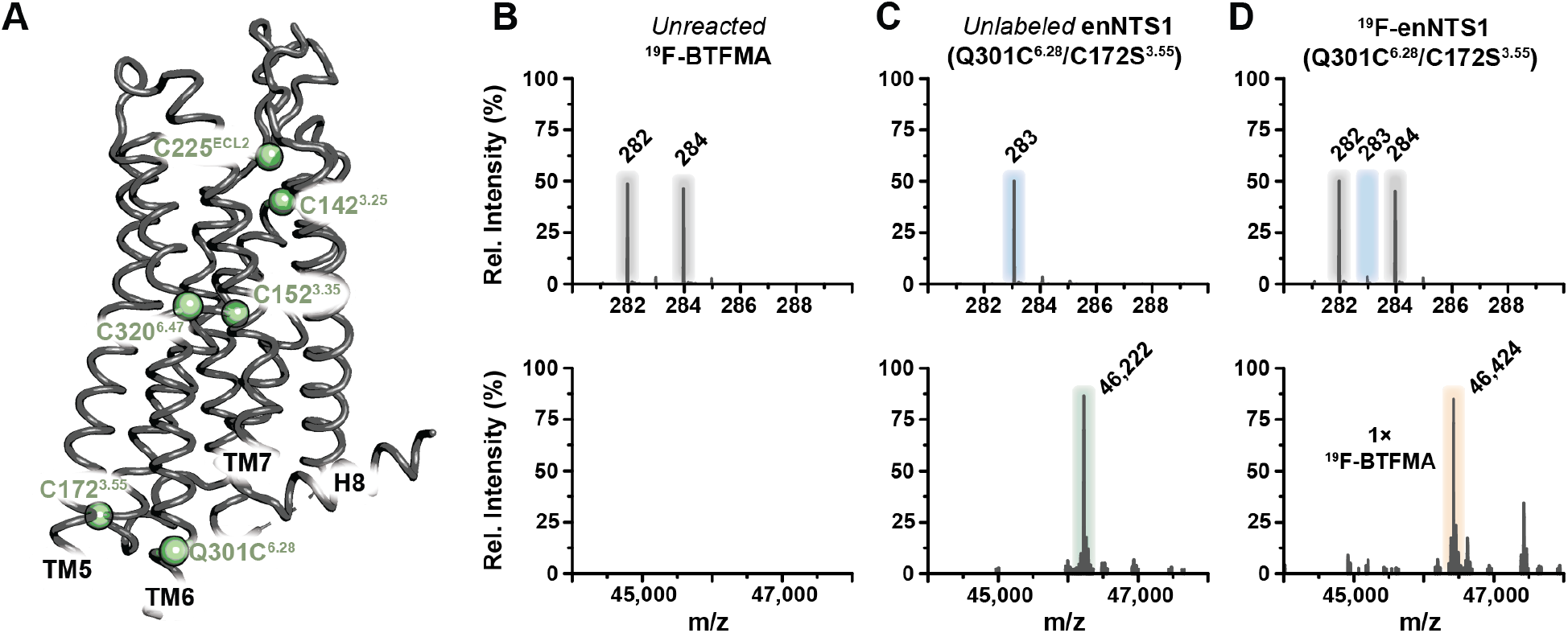
Detergent micelles sequester ^19^F-BTFMA probe molecules. (A) Atomic model of enNTS1 (PDB 4BWB) highlighting cysteine residue mutations (green spheres).^37^ LC-MS results of (B) detergent-solubilized ^19^F-BTFMA, (C) unlabeled enNTS1(Q301C^6.28^/C172S^3.55^), and (D) ^19^F-enNTS1(Q301C^6.28^/C172S^3.55^). All LC-MS peak intensities are relative to each individual spectrum. ^19^F-BTFMA m/z = 282/284 Da; enNTS1(Q301C^6.28^/C172S^3.55^) m/z = 46,222 Da; ^19^F-enNTS1(Q301C^6.28^/C172S^3.55^) m/z = 46,424 Da (+1 ^19^F-BTFMA molecule). A MW intensity of 283 Da was also observed in all enNTS1 protein samples, regardless of ^19^F-BTFMA presence, corresponding to an unrelated sample contaminate.

Next, we collected a spectrum of unreacted ^19^F-BTFMA under identical buffer conditions [Fig. 1(H)]. It contained the same two resonances at 13.6 and 14.8 ppm, but with narrower linewidths than the ^19^F-enNTS1 samples. We assigned these two ^19^F resonances, using ^1^H 1D spectrum, as ^19^F-BTFMA and a triethylamine trifluoride reactant impurity [Fig. S1]. Given that unreacted ^19^F-BTFMA is considerably lipophilic with a theoretical octanol:water partition coefficient (logP) ∼3.5,^29^ we hypothesized that detergent micelles may be sequestering excess ^19^F-probe molecules. We turned to liquid-chromatography mass spectrometry (LC-MS) to test this hypothesis. The reverse-phase LC step separates all non-covalent components of the proteomicelle for accurate determination of individual molecular weights. ^19^F-BTFMA solubilized in detergent micelles showed the expected 282 and 284 Da Bromine isotope doublet pattern of the protonated, unreacted form [Fig. 2(B)]. Unlabeled enNTS1(Q301C^6.28^/C172S^3.55^) exhibited a prominent intensity of 46,222 Da [Fig. 2(C)]. LC-MS analysis of ^19^F-enNTS1(Q301C^6.28^/C172S^3.55^) contained a major intensity of 46,424 Da, corresponding to conjugation of a single ^19^F-BTFMA molecule (+202 Da), as well as the 282 and 284 Da doublet of unreacted ^19^F-BTFMA [Fig. 2(D)]. Thus, the LC-MS illustrates that detergent micelles can sequester unreacted ^19^F-BTFMA.

We hypothesized that removal of excess, unreacted ^19^F-BTFMA molecules prior to detergent solubilization would eliminate micellar sequestration. We sonicated cell pellets containing enNTS1(Q301^6.28^/C172S^3.55^), incubated for 1 h with ^19^F-BTFMA, and then performed a membrane preparation via ultracentrifugation. The sample was pelleted at 100k g, decanted, and washed with buffer containing no detergent. This ultracentrifugation step was repeated to insure complete removal of excess ^19^F-probe. The ^19^F-enNTS1(Q301^6.28^/C172S^3.55^) sample was then solubilized with DM detergent and purified as above to produce a ^19^F-NMR spectrum with no observable signals [Fig. 3(A)]. Applying this methodology to ^19^F-enNTS1(Q301C^6.28^/C172S^3.55^) yielded a spectrum containing a single resonance in slow exchange [Fig. 3(B)].^16^ Intact and protease-digested LC-MS confirmed that ^19^F-enNTS1(Q301C^6.28^/C172S^3.55^) was exclusively-labeled at Q301C^6.28^ with no observable unreacted ^19^F-BTFMA [Fig. 3(C) and Table S1]. Thus, we propose a simple four-step protocol for Selective Labeling Absent of Probe Sequestration (SLAPS): physically-disrupt cell membranes in the absence of detergent, incubate membranes with cysteine-reactive ^19^F-probes, remove excess unreacted ^19^F-probe molecules via ultracentrifugation, and finally solubilize in detergent of choice [Fig. 4]. Without consideration of probe sequestration by detergent micelles as a source of offsite labeling, spectral deconvolution of ^19^F-enNTS1(Q301C^6.28^/C172S^3.55^) prepared using conventional methods vs SLAPS would both be compatible with at least three receptor conformations in slow exchange [Fig. 3(D)].

**Figure 3.**
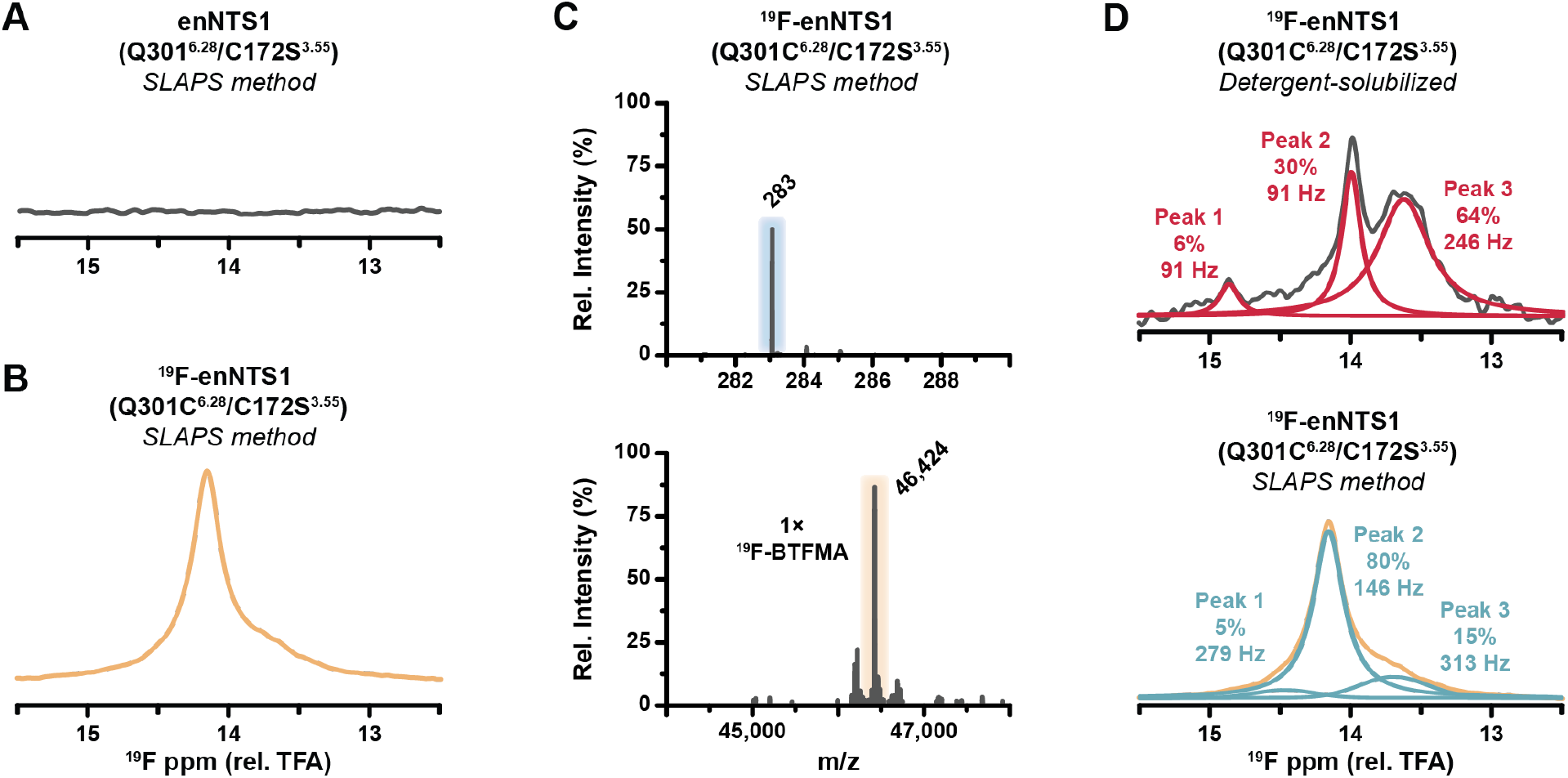
Selective Labeling Absent of Probe Sequestration (SLAPS) eliminates offsite reactions in enNTS1. ^19^F-NMR spectra of (A) enNTS1(Q301^6.28^/C172S^3.55^) and (B) ^19^F-enNTS1(Q301C^6.28^/C172S^3.55^) prepared using the SLAPS protocol. Note that ^19^F-BTFMA only reacts with enNTS1(Q301C^6.28^/C172S^3.55^). (C) LC-MS spectra of ^19^F-enNTS1(Q301C^6.28^/C172S^3.55^) purified using SLAPS. Note the absence of unreacted ^19^F-BTFMA (top). (D) MestReNova^32^ spectral deconvolution of ^19^F-enNTS1(Q301C^6.28^/C172S^3.55^) labeled while solubilized in detergent micelles (top) or using the SLAPS methodology (bottom). Final sample buffer conditions for all NMR spectra: 20 mM HEPES, 50 mM NaCl, 50 µM TFA, and 0.01% (w/v) LMNG at pH 7.5. All NMR samples were supplemented with 10% (v/v) D_2_O.

**Figure 4.**
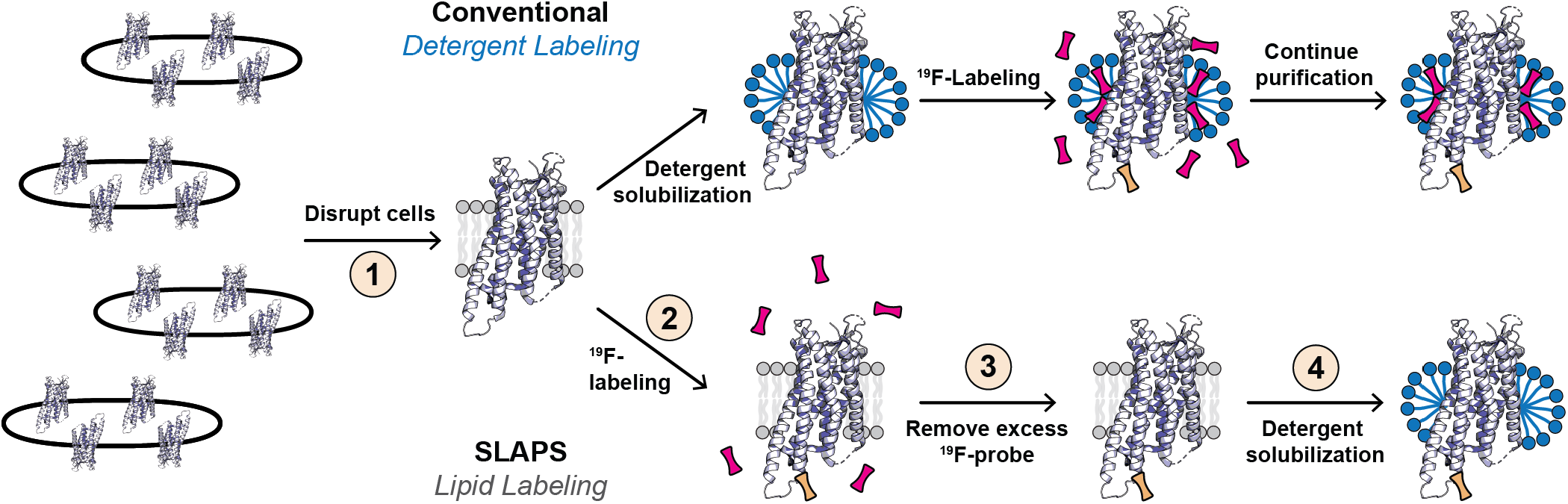
Selective Labeling Absent of Probe Sequestration (SLAPS) protocol. (top) Model illustrating that receptors which are solubilized in detergent, prior to ^19^F-labeling, will sequester unreacted probe molecules (red) in addition to the correctly conjugated probe (orange). SLAPS follows a simple four-step protocol: 1) physically-disrupt cell membranes in the absence of detergent, 2) incubate membranes with cysteine-reactive ^19^F-probes, 3) remove excess unreacted ^19^F-probe molecules via ultracentrifugation, and 4) solubilize in the detergent of choice.

## Conclusions

Conjugation of trifluoromethyl probes to GPCRs solubilized in detergent micelles is known to results in offsite cysteine reactions, but undesired reactions can be eliminated using the IMCM approach.^19^ Here, we characterize a second source of offsite labeling: non-covalent probe sequestration by detergent micelles. Although IMCM appears less effective at eliminating probe sequestration, there are several important considerations in our application of IMCM to enNTS1. First, the IMCM method was originally developed for the 2,2,2-triflouroethyl-1-thiol (^19^F-TET) probe, which is considerably less lipophilic (logP ∼1.5) than ^19^F-BTFMA.^29^ Second, the ^19^F-labeled receptors used in that study were solubilized in n-dodecyl-β-D-maltopyranoside (DDM), whose micelles retain receptor structural integrity better than the DM detergent used for enNTS1.^30^ Thus, we hypothesize that the IMCM approach would also limit detergent sequestration under these specific conditions. Although, it may be less effective with the numerous more lipophilic probes such as 3-bromo-1,1,-trifluoroproan-2one (BTFA; logP ∼2.0), 1-bromo-3,3,4,4,4-pentafluorobutan-2one (BPFB; logP ∼2.7), and N-(4-bromo-3-(trifluoromethyl)phenyl)acetamide (3-BTFMA; logP ∼2.9).^22^ SLAPS should be broadly applicable to a variety of cysteine-reactive lipophilic probes, other GPCRs, and additional membrane protein classes solubilized in detergent micelles or lipid mimetics.

## Materials and Methods

### enNTS1 plasmid construct and protein expression

The previously characterized functional variant enNTS_1_^27^ was available in an expression vector (termed pDS170) with an open reading frame encoding an N-terminal maltose-binding protein signal sequence (MBPss), followed by a 10x His tag, a maltose-binding protein (MBP), a NNNNNNNNNNG linker and a HRV 3C protease site (LEVLFQGP) which were linked via a BamHI restriction site (resulting in additional residues GS) to residue T42 of the receptor. C-terminally T416 of the receptor was linked via a NheI restriction site (resulting in additional residues AS) to an Avi-tag for in vivo biotinylation, a HRV 3C protease site, a GGSGGS linker and a monomeric ultra-stable green fluorescent protein (muGFP).^31^ enNTS1 plasmids were transformed into BL21(DE3) *Escherichia coli* cells and plated overnight on LB agar supplemented with 100 µg/mL carbenicillin at 37 °C. Liquid LB starter cultures were supplemented with 100 µg/mL carbenicillin, seeded with colonies, and incubated overnight at 37 °C and 220 RPM. One-liter 2xYT media supplemented with 100 µg/mL carbenicillin and 0.3% (w/v) glucose were inoculated with overnight LB starter culture, and incubated at 37 °C and 220 RPM to an OD_600_ ≅ 0.15. The cultures were then cooled to 16 °C. Once each culture reached an OD_600_ ≅ 0.6, they were induced with 0.3 mM IPTG and incubated for ∼ 21h at 16 °C and 220 RPM. The cultures were harvested via centrifugation at 4,000 g and stored at -80 °C.

### Selective Labeling Absent of Probe Sequestration (SLAPS) protocol

Cell pellets were solubilized on ice in *solubilization buffer* (100 mM HEPES, 400 mM NaCl, 20% (v/v) glycerol, 10 mM MgCl_2_, 10 mM imidazole, pH 8.0), 100 mg lysozyme, 1-unit DNAse, 0.2 mM PMSF, and one protease inhibitor cocktail tablet. Solution was then sonicated on ice: two minutes processing time (10 s on, 20 s off) at 30% maximum amplitude. Following sonication, 5 mM ^19^F-BTFMA and 0.2 mM PMSF was added to the solution and stirred at 4 °C for one hour. After incubation, 16 mg aldrithiol was added to the solution and stirred at 4 °C for an additional 10 minutes. To remove excess ^19^F-BTFMA probe, a membrane preparation was performed via ultracentrifugation at 100,000 g for 10 minutes. The supernatant was decanted and the pellet was resuspended in the same volume of fresh *solubilization buffer*. A second membrane preparation was performed via ultracentrifugation at 100,000 g for 10 minutes.

### enNTS1 protein purification

Following ultracentrifugation, the receptor sample was solubilized at a final concentration of 0.6% (w/v) CHAPS, 0.12% (w/v) CHS, and 1% (w/v) DM detergent. The solution was stirred at 4 °C for two hours. After incubation, native lipids were removed via centrifugation at 100,000 g for 10 minutes. The remaining enNTS1 supernatant was then incubated with *equilibrated TALON* resin (25 mM HEPES, 10% (v/v) glycerol, 300 mM NaCl, 0.15% (w/v) DM, pH 8.00) at 4 °C for 15 minutes. Following TALON resin binding, the receptor solution was placed into a gravity column to remove unbound proteins. The TALON resin was then subjected to two subsequent wash steps: *TALON wash #1* (25 mM HEPES, 10% (v/v) glycerol, 500 mM NaCl, 0.15% (w/v) DM, 10 mM Imidazole, 4 mM ATP, 10 mM MgCl2, pH 8.0) and *TALON wash #2* (25 mM HEPES, 10% (v/v) glycerol, 350 mM NaCl, 0.1% (w/v) LMNG, 10 mM Imidazole, pH 8.0). It is important to note that the second wash step also serves as a detergent exchange step from DM to LMNG. Following detergent exchange, enNTS1 was eluted with *TALON elute buffer* (25 mM HEPES, 10% (v/v) glycerol, 500 mM NaCl, 0.01% (w/v) LMNG, 350 mM Imidazole, pH 8.0) and incubated with 3 mg 3C precision protease for 2-16 h at 4 °C to remove MBP and muGFP expression tags. The cleaved enNTS1 was concentrated in a 50 MWCO concentrator via centrifugation at 3,500 g and then diluted 10-fold in *SP equilibration buffer* (20 mM HEPES, 10% (v/v) glycerol, 0.01% (w/v) LMNG, pH 7.4) was added. This resulting solution was loaded onto an equilibrated 5 mL SP ion-exchange (IEX) column via GE AKTA Pure system. The SP IEX column was washed with *SP wash buffer* (20 mM HEPES, 10% (v/v) glycerol, 250 mM NaCl, 0.01% (w/v) LMNG, pH 7.4) until AU_280_ was stable. An equilibrated 1 mL Ni^2+^-NTA column was attached in-tandem following the 5 mL SP IEX column, and the receptor eluted with *SP elute buffer* (20 mM HEPES, 10% (v/v) glycerol, 1 M NaCl, 0.01% (w/v) LMNG, 25 mM Imidazole, pH 7.4). The enNTS1 solution was then concentrated in a 50 MWCO concentrator via centrifugation at 3,500 g and injected onto a GE S200 Increase SEC column equilibrated in *NMR buffer* (20 mM HEPES, 50 mM NaCl, 0.01% (w/v) LMNG, 50 µM TFA, pH 7.5). Following SEC, the desired enNTS1 fractions were pooled, concentrated to 100-300 µM, and flash-frozen via liquid nitrogen and stored at -80 °C.

### ^19^F NMR

^19^F NMR spectra were collected on a 14.1 T Bruker AVANCE NEO (Indiana University – Bloomington) spectrometer equipped with a 5mm TCI CryoProbe tunable to the fluorine frequency. Free induction decay (FID) signals were collected by applying a π/2 pulse length of 13.5 µs, a recycling time of 0.8 ms, and an acquisition time of 0.15 s. A total of 8192 scans were collected generating a FID comprised of 2499 complex points which were zero-filled to 8000 complex points, and apodized with a 30 Hz exponential filter. NMR spectra were deconvoluted using MestReNova as previously detailed.^16,32^

### Mass spectrometry

Intact protein analysis - Samples were analyzed on a Synapt G2S equipped with an iClass Acquity HPLC (Waters). Buffer A was 0.1% (v/v) formic acid in water and Buffer B was 0.1% (v/v) formic acid in acetonitrile. Proteins were separated using a nine-minute gradient from 5-99% Buffer B at a flow rate of 50 nL/min. Proteins were separated using a 5 cm x 0.5 mm column in-house packed with Jupiter 5μm C4 resin (Phenomenex). The ToF was set to scan at 1 s intervals and an analyzer setting of “Resolution” was used. Protein processing -Samples were resuspended and denatured in 8 M urea with 100 mM ammonium bicarbonate (pH 7.8). Disulfide bonds were reduced by incubation for 45 min at 57 °C with a final concentration of 10 mM Tris (2-carboxyethyl) phosphine hydrochloride (#C4706, Sigma Aldrich). A final concentration of 20 mM iodoacetamide (#I6125, Sigma Aldrich) was then added to alkylate these side chains and the reaction was allowed to proceed for one hour in the dark at 21 °C. Samples were diluted to 1 M urea using 100 mM ammonium bicarbonate, pH 7.8. Trypsin (V5113, Promega) or chymotrypsin (#11418467001, Sigma Aldrich) was added at a 1:100 ratio and the samples were digested for 14 hours at 37 °C. Mass spectrometry - individual samples were desalted using ZipTip pipette tips (EMD Millipore), dried down and resuspended in 0.1% (v/v) formic acid. Fractions were analyzed by LC-MS on an Orbitrap Fusion Lumos equipped with an Easy NanoLC1200 HPLC (Thermo Fisher Scientific). Buffer A was 0.1% (v/v) formic acid in water. Buffer B was 0.1% (v/v) formic acid in 80% acetonitrile. Peptides were separated on a 30-minute gradient from 0-3% Buffer B. Precursor ions were measured in the Orbitrap with a resolution of 120,000. Fragment ions were measured in the Orbitrap with a resolution of 15,000. The spray voltage was set at 1.8 kV. Orbitrap MS1 spectra (AGC 1×10^6^) were acquired from 350-2000 m/z followed by data-dependent HCD MS/MS (collision energy 30%, isolation window of 2 Da) for a 3 s cycle time. Charge state screening was enabled to reject unassigned and singly charged ions. A dynamic exclusion time of 30 s was used to discriminate against previously selected ions. Database search - The LC-MS/MS data was searched against the protein sequence using Protein Prospector (v5.22.1). The database search parameters for the tryptic search allowed for two missed cleavages and one non-tryptic cleavage. The search parameters for the chymotryptic search allowed for four missed cleavages and one non-chymotryptic cleavage. A precursor and fragment mass tolerance of 10 ppm was used. Oxidation of methionine, pyroglutamine on peptide amino termini, carbamidomethylation of cysteine, and protein N-terminal acetylation were set as variable modifications. In addition, modification of cysteine residues by conjugated ^19^F-BTFMA (C_9_H_6_F_3_NO) was set as a variable modification.

## Supplementary Material

^1^H-NMR spectrum of ^19^F-BTFMA probe and protease digestion LC-MS result of ^19^F-enNTS1(Q301C^6.28^/C172S^3.55^).

## Acknowledgments

We are grateful to Dr. Hongwei Wu at Indiana University for NMR instrument assistance and Prof. Daniel Scott at the Florey Institute for providing the enNTS1 plasmid used in this study. We appreciate the constructive feedback from James Bower, Dr. Scott Robson, Thomas Shriver, and Skylar Zemmer. The project was funded by: Indiana Precision Health Initiative (JJZ), NIH grants R00GM115814 (JJZ) and R35GM143054 (JJZ). The 14.1 T spectrometer used in this study was generously supported by the Indiana University Fund.

## Supporting Information

**Figure S1.**
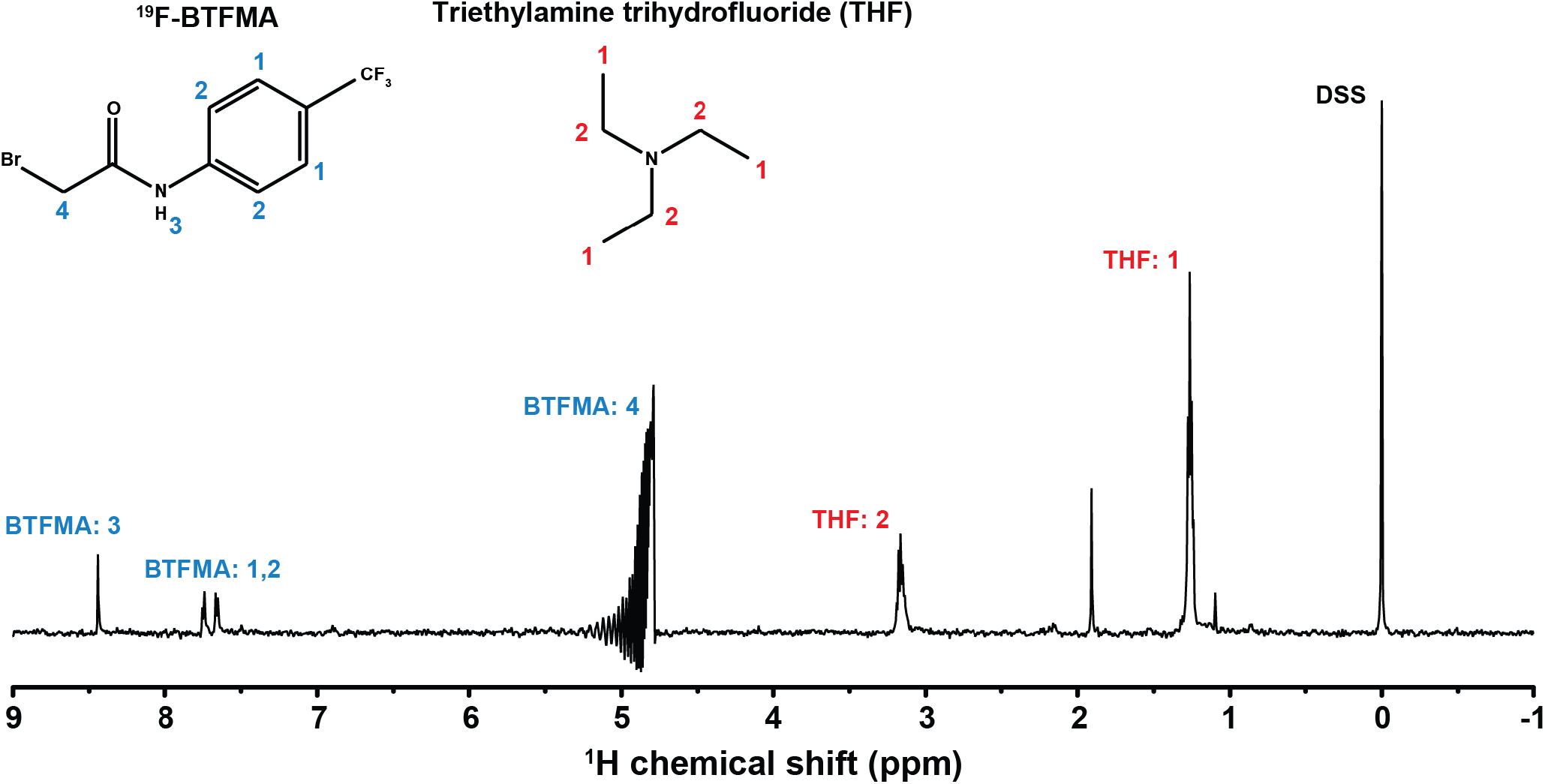
^1^H NMR spectrum of ^19^F-BTFMA probe. ^19^F-BTFMA probe (100 µM) solubilized in H_2_O. DSS standard (20 µM) was used as a reference. Small molecule triethylamine trihydrofluoride (THF) is observed as a contaminating reagent from ^19^F-BTFMA synthesis. Spectrum was collected in a 3 mm O.D. tube. Free induction decay (FID) signals were collected by applying a π/2 pulse length of 7.25 μs with an acquisition time of 0.72 s. A total of 64 scans were collected to generate an FID comprised of 5,120 complex points which were zero-filled to 8,000 complex points, and processed with a cosine-squared window function.

**Table S1.**
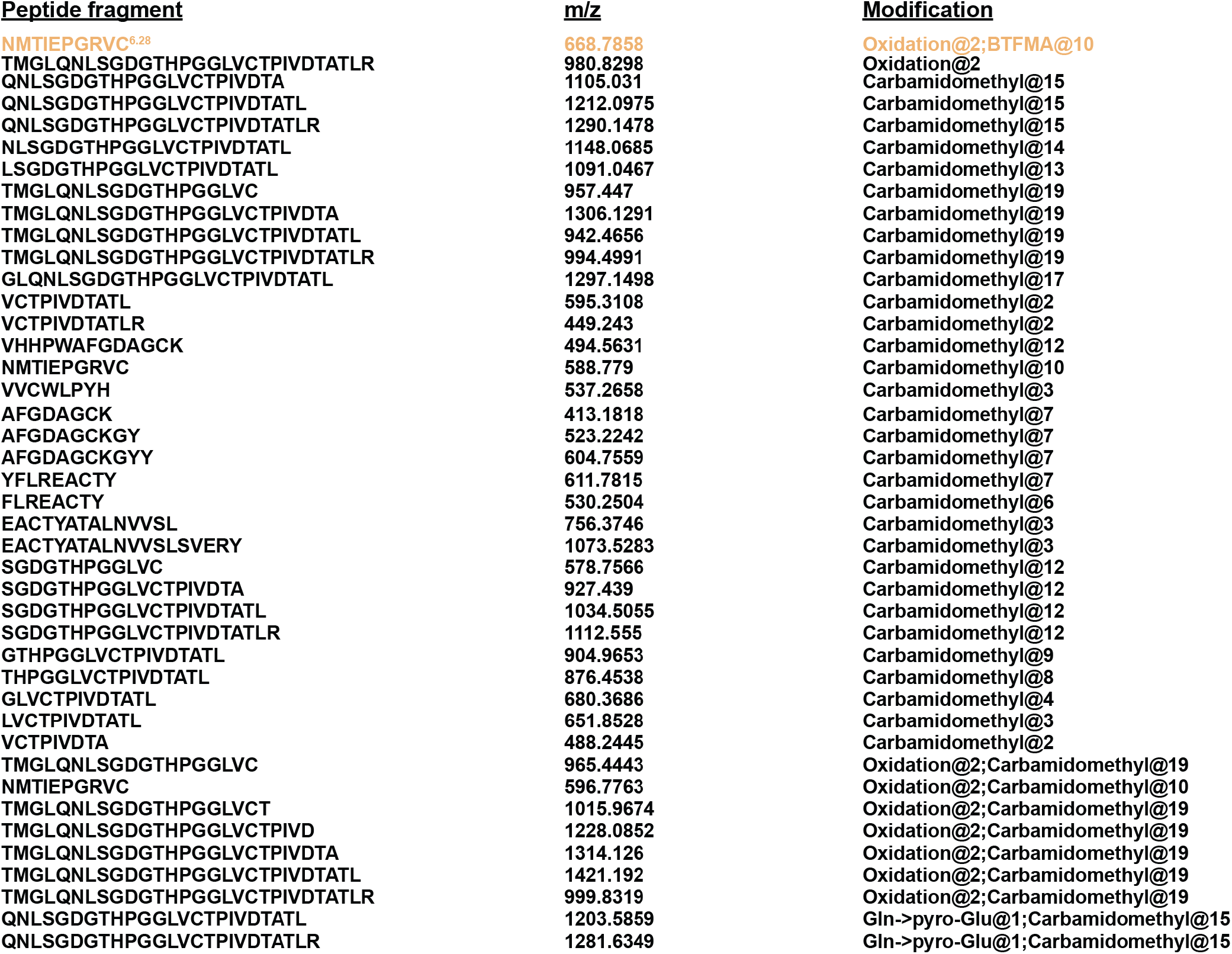
Database search for LC-MS/MS of protease-digested ^19^F-enNTS1(Q301C^6.28^/C172S^3.55^) labeled using SLAP approach. ^19^F-enNTS1(Q301C^6.28^/C172S^3.55^) was digested with trypsin and chymotrypsin then analyzed by LC-MS/MS. Peptide fragments of covering all five remaining cysteine residues were detected. Mass-to-charge (m/z) ratio and residue modification are listed for each peptide fragment. Note only C301^6.28^ (Peach) incorporates ^19^F-BTFMA during SLAPS labeling.

## References

1. de Mendoza A, Sebé-Pedrós A, Ruiz-Trillo I (2014) The evolution of the GPCR signaling system in eukaryotes: modularity, conservation, and the transition to metazoan multicellularity. Genome Biol. Evol. 6:606–619.

2. Hanlon CD, Andrew DJ (2015) Outside-in signaling – a brief review of GPCR signaling with a focus on the Drosophila GPCR family. J. Cell Sci. 128:3533–3542.

3. Schönegge A-M, Gallion J, Picard LP, Wilkins AD, Le Gouill C, Audet M, Stallaert W, Lohse MJ, Kimmel M, Lichtarge O, et al. (2017) Evolutionary action and structural basis of the allosteric switch controlling β2AR functional selectivity. Nat. Commun. 8:2169.

4. Hauser AS, Attwood MM, Rask-Andersen M, Schiöth HB, Gloriam DE (2017) Trends in GPCR drug discovery: new agents, targets and indications. Nat. Rev. Drug Discov. 16:829–842.

5. Lu S, He X, Yang Z, Chai Z, Zhou S, Wang J, Rehman AU, Ni D, Pu J, Sun J, et al. (2021) Activation pathway of a G protein-coupled receptor uncovers conformational intermediates as targets for allosteric drug design. Nat. Commun. 12:4721.

6. Zhou Y, Meng J, Xu C, Liu J (2021) Multiple GPCR Functional Assays Based on Resonance Energy Transfer Sensors. Front. Cell Dev. Biol. 9.

7. Ye S, Zaitseva E, Caltabiano G, Schertler GFX, Sakmar TP, Deupi X, Vogel R (2010) Tracking G-protein-coupled receptor activation using genetically encoded infrared probes. Nature 464:1386–1389.

8. Manglik A, Kim TH, Masureel M, Altenbach C, Yang Z, Hilger D, Lerch MT, Kobilka TS, Thian FS, Hubbell WL, et al. (2015) Structural Insights into the Dynamic Process of β2-Adrenergic Receptor Signaling. Cell 161:1101–1111.

9. Kim TH, Chung KY, Manglik A, Hansen AL, Dror RO, Mildorf TJ, Shaw DE, Kobilka BK, Prosser RS (2013) The Role of Ligands on the Equilibria Between Functional States of a G Protein-Coupled Receptor. J. Am. Chem. Soc. 135:9465–9474.

10. Verardi R, Traaseth NJ, Masterson LR, Vostrikov V V, Veglia G (2012) Isotope labeling for solution and solid-state NMR spectroscopy of membrane proteins. Adv. Exp. Med. Biol. 992:35–62.

11. Frei JN, Broadhurst RW, Bostock MJ, Solt A, Jones AJY, Gabriel F, Tandale A, Shrestha B, Nietlispach D (2020) Conformational plasticity of ligand-bound and ternary GPCR complexes studied by 19F NMR of the β1-adrenergic receptor. Nat. Commun. 11:669.

12. Liu JJ, Horst R, Katritch V, Stevens RC, Wüthrich K (2012) Biased Signaling Pathways in β2-Adrenergic Receptor Characterized by 19F-NMR. Science (80). 335:1106 LP – 1110.

13. Sušac L, Eddy MT, Didenko T, Stevens RC, Wüthrich K (2018) A2A adenosine receptor functional states characterized by 19F-NMR. Proc. Natl. Acad. Sci. 115:12733 LP – 12738.

14. Klein-Seetharaman J, Getmanova EV, Loewen MC, Reeves PJ, Khorana HG (1999) NMR spectroscopy in studies of light-induced structural changes in mammalian rhodopsin: Applicability of solution 19F NMR. Proc. Natl. Acad. Sci. 96:13744–13749.

15. Wang X, Liu D, Shen L, Li F, Li Y, Yang L, Xu T, Tao H, Yao D, Wu L, et al. (2021) A Genetically Encoded F-19 NMR Probe Reveals the Allosteric Modulation Mechanism of Cannabinoid Receptor 1. J. Am. Chem. Soc. 143:16320–16325.

16. Dixon AD, Inoue A, Robson SA, Culhane KJ, Trinidad JC, Sivaramakrishnan S, Bumbak F, Ziarek JJ (2022) Effect of Ligands and Transducers on the Neurotensin Receptor 1 Conformational Ensemble. J. Am. Chem. Soc. 144:10241–10250.

17. Chen H, Viel S, Ziarelli F, Peng L (2013) 19F NMR: a valuable tool for studying biological events. Chem. Soc. Rev. 42:7971–7982.

18. Staus DP, Wingler LM, Pichugin D, Prosser RS, Lefkowitz RJ (2019) Detergent-and phospholipid-based reconstitution systems have differential effects on constitutive activity of G-protein-coupled receptors. J. Biol. Chem. 294:13218–13223.

19. Sušac L, O’Connor C, Stevens RC, Wüthrich K (2015) In-Membrane Chemical Modification (IMCM) for Site-Specific Chromophore Labeling of GPCRs. Angew. Chem. Int. Ed. Engl. 54:15246–15249.

20. Klein-Seetharaman J, Oikawa M, Grimshaw SB, Wirmer J, Duchardt E, Ueda T, Imoto T, Smith LJ, Dobson CM, Schwalbe H (2002) Long-range interactions within a nonnative protein. Science 295:1719–1722.

21. Luchette PA, Prosser RS, Sanders CR (2002) Oxygen as a paramagnetic probe of membrane protein structure by cysteine mutagenesis and (19)F NMR spectroscopy. J. Am. Chem. Soc. 124:1778–1781.

22. Ye L, Larda ST, Frank Li YF, Manglik A, Prosser RS (2015) A comparison of chemical shift sensitivity of trifluoromethyl tags: optimizing resolution in <sup>19</sup>F NMR studies of proteins. J. Biomol. NMR 62:97–103.

23. Mulry E, Ray AP, Eddy MT (2021) Production of a Human Histamine Receptor for NMR Spectroscopy in Aqueous Solutions. Biomolecules 11.

24. Eddy MT, Didenko T, Stevens RC, Wüthrich K (2016) β2-Adrenergic Receptor Conformational Response to Fusion Protein in the Third Intracellular Loop. Structure 24:2190–2197.

25. Liu JJ, Horst R, Katritch V, Stevens RC, Wüthrich K (2012) Biased signaling pathways in β2-adrenergic receptor characterized by 19F-NMR. Science 335:1106–1110.

26. Sanchez-Soto M, Verma RK, Willette BKA, Gonye EC, Moore AM, Moritz AE, Boateng CA, Yano H, Free RB, Shi L, et al. (2020) A structural basis for how ligand binding site changes can allosterically regulate GPCR signaling and engender functional selectivity. Sci. Signal. 13.

27. Bumbak F, Keen AC, Gunn NJ, Gooley PR, Bathgate RAD, Scott DJ (2018) Optimization and 13CH3 methionine labeling of a signaling competent neurotensin receptor 1 variant for NMR studies. Biochim. Biophys. Acta - Biomembr. 1860:1372–1383.

28. Ballesteros JA, Weinstein H (1995) Integrated methods for the construction of three-dimensional models and computational probing of structure-function relations in G protein-coupled receptors. Receptor Molecular Biology. Vol. 25. Academic Press; 1995. pp. 366–428.

29. Anon MarvinSketch was used for calculating logP partition coefficients, MarvinSketch version 20.17.0, ChemAxon (https://www.chemaxon.com).

30. Lee S, Mao A, Bhattacharya S, Robertson N, Grisshammer R, Tate CG, Vaidehi N (2016) How Do Short Chain Nonionic Detergents Destabilize G-Protein-Coupled Receptors? J. Am. Chem. Soc. 138:15425–15433.

31. Scott DJ, Gunn NJ, Yong KJ, Wimmer VC, Veldhuis NA, Challis LM, Haidar M, Petrou S, Bathgate RAD, Griffin MDW (2018) A Novel Ultra-Stable, Monomeric Green Fluorescent Protein For Direct Volumetric Imaging of Whole Organs Using CLARITY. Sci. Rep. 8:667.

32. Willcott MR (2009) MestRe Nova. J. Am. Chem. Soc. 131:13180.

33. Miller-Gallacher JL, Nehmé R, Warne T, Edwards PC, Schertler GFX, Leslie AGW, Tate CG (2014) The 2.1 Å Resolution Structure of Cyanopindolol-Bound β1-Adrenoceptor Identifies an Intramembrane Na+ Ion that Stabilises the Ligand-Free Receptor. PLoS One 9:e92727.

34. Vadim C, Rosenbaum DM, Hanson MA, Rasmussen SGF, Thian FS, Kobilka TS, Choi HJ, Kuhn P, Weis WI, Kobilka KB, Stevens RC (2007) High-Resolution Crystal Structure of an Engineered Human β2-Adrenergic G Protein–Coupled Receptor. Science (80). 318:1258–1265.

35. Wei L, Chun E, Thompson AA, Chubukov P, Xu F, Katritch V, Han GW, Roth CB, Heitman LH, Ijzerman AP, Cherezov P, Stevens RC (2012) Structural Basis for Allosteric Regulation of GPCRs by Sodium Ions. Science (80). 337:232–236.

36. Okada T, Sugihara M, Bondar A-N, Elstner M, Entel P, Buss V (2004) The Retinal Conformation and its Environment in Rhodopsin in Light of a New 2.2Å Crystal Structure. J. Mol. Biol. 342:571–583.

37. Egloff P, Hillenbrand M, Klenk C, Batyuk A, Heine P, Balada S, Schlinkmann KM, Scott DJ, Schütz M, Plückthun A (2014) Structure of signaling-competent neurotensin receptor 1 obtained by directed evolution in Escherichia coli. Proc. Natl. Acad. Sci. 111:E655 LP–E662.

